# mafsmith: a Rust reimplementation of vcf2maf

**DOI:** 10.64898/2026.05.12.724685

**Authors:** Robert Allaway

## Abstract

The Mutation Annotation Format (MAF) is a standard interchange format for somatic variant data in tumor genomics. Converting variant call format (VCF) files to MAF requires functional annotation (through tools such as the Ensembl Variant Effect Predictor) and complex allele normalisation and field-mapping logic. The gold-standard implementation, vcf2maf, is written in Perl and could be made more computationally efficient by translating it to a newer language and adding support for parallel processing. Here we describe mafsmith, an implementation of vcf2maf in Rust. The mafsmith implementation of vcf2maf reimplements the allele-normalisation and field-mapping logic of vcf2maf and uses fastVEP for annotation, achieving field-for-field identical output across fifteen validated caller types and formats spanning germline, somatic, structural variant, and annotation-database VCFs. When both tools are run with the same Ensembl VEP annotation cache, mafsmith `vcf2maf` produces 0 conversion differences versus vcf2maf across 23 diverse datasets aligned to GRCh38 or GRCh37. The companion `maf2vcf`, `vcf2vcf`, and `maf2maf` subcommands were similarly validated against their reference Perl counterparts across six datasets. Benchmarked on multiple reference samples totalling 27.5 million variants, mafsmith achieves approximately 80-fold faster conversion of pre-annotated VCFs (range 74.3–84.1×), enabling faster and cheaper conversion of vcfs to mafs. mafsmith is open source under the same license as vcf2maf and available at https://github.com/nf-osi/mafsmith.

## Introduction

Somatic variant calling produces VCF files whose downstream use in cancer genomics almost universally requires conversion to the Mutation Annotation Format (MAF). MAF is a primary data format for the NCI Genomic Data Commons (GDC)^1^, which hosts large cancer cohorts including The Cancer Genome Atlas^2^, and is consumed by widely used analytical tools including maftools^3^, cBioPortal^4^, and OncoKB^5^. The conversion from VCF to MAF is non-trivial: it requires functional annotation of each variant against a transcript database, selection of a canonical or preferred transcript, normalisation of multi-allelic and indel representations, and mapping of genotype information to allele-level MAF fields.

The standard tool for this conversion is vcf2maf^6^. vcf2maf is a Perl script that wraps the Ensembl Variant Effect Predictor (VEP)^7^, handling the full complexity of VCF allele representations, multi-allelic sites, structural variants, and caller-specific FORMAT field conventions that have accumulated over years of production use across major cancer genomics consortia. Its breadth of supported vcf versions and spec-non-conformant files make it the reference implementation.

However, vcf2maf has performance limitations, described further in the results section. For large cohorts of hundreds or thousands of samples, this becomes a major computational burden. The dependency on a compatible VEP and reference database can also create significant installation challenges. Recent development of fastVEP^8^, a reimplementation of VEP’s core annotation logic in the Rust programming language, substantially reduces annotation time, achieving up to 130-fold speedup over the original Perl VEP while maintaining complete concordance. However, the conversion step itself (allele normalisation, genotype parsing, field mapping) still requires vcf2maf.

Here we describe mafsmith, a complete rewrite of vcf2maf in Rust that pairs naturally with fastVEP to replace the entire vcf2maf + VEP stack with a single, self-contained toolchain. Both tools leverage Rust’s performance characteristics and native parallelism to achieve throughput that is not possible with the standard implementations. mafsmith reimplements the allele-normalisation and field-mapping logic of vcf2maf from first principles, targeting field-for-field identical output. We validate all four conversion subcommands (`vcf2maf`, `maf2vcf`, `vcf2vcf`, and `maf2maf`) against their reference Perl counterparts, demonstrate 79.4-fold speedup for the `vcf2maf` conversion step, and describe the specific edge cases and caller-specific conventions that required careful evaluation to achieve full concordance.

## Methods

### Development approach

mafsmith was implemented in Rust using Anthropic’s Claude Sonnet 4.6 language model assisted by the Claude Code command-line interface. Development followed an iterative cycle of implementation and validation: after each implementation pass, mafsmith output was compared field-by-field to vcf2maf output on real-world VCFs representing a diverse panel of sequencing platforms, variant callers, and VCF conventions. Discrepancies were diagnosed to their root cause and resolved in targeted fixes before the next validation cycle. The datasets listed in Table 1 and described in the next section therefore represent both the performance evaluation corpora and the data against which the conversion logic was defined and refined.

**Table 1.** Validation of mafsmith against vcf2maf (--strict mode; 0 conversion-field mismatches in all cases; 87.2 million variants compared in total).

| Caller | Source | Variants compared |
| --- | --- | --- |
| DeepVariant 1.2.0 | syn31624545<sup>28</sup> | 2,936,426 |
| GATK MuTect2 (single-sample) | syn31624525<sup>28</sup> | 751,548 |
| GATK MuTect2 (paired T/N) | GIAB HG008<sup>12</sup>; SEQC2 HCC1395<sup>29</sup> | 549,590 variants from 2 samples |
| FreeBayes | syn31624535<sup>28</sup> | 2,858,303 |
| Strelka2 germline | syn31624939; syn31624637<sup>28</sup>; syn29349669<sup>30</sup> | 9,327,717 variants from 2 samples |
| Strelka2 somatic SNVs | GIAB HG008<sup>12</sup>; SEQC2 HCC1395<sup>29</sup> | 3,754,567 variants from 2 samples |
| Strelka2 somatic indels | syn68172710<sup>31</sup>; GIAB HG008<sup>12</sup> | 317,094 variants from 2 samples |
| SV callers (Manta / DELLY) | syn21296193, syn21296175, syn21296184<sup>32</sup> | 398 (all SVs) |
| VarScan2 somatic | syn6840402<sup>31</sup> | 59,618 |
| VarDict (paired T/N) | syn6039268<sup>31</sup> | 9,303,064 |
| SomaticSniper | SEQC2 HCC1395<sup>29</sup> | 164,704 |
| GIAB germline benchmarks | HG001-HG007 (NIST v4.2.1)<sup>12</sup> | 27,529,001 variants from 7 samples |
| ICGC PCAWG consensus (SNV/MNV) | ICGC PCAWG open data (GRCh37)<sup>20</sup> | 21,628,933 variants from 1,902 samples |
| DepMap CCLE WGS (GATK MuTect2) | DepMap CCLE (hg38)<sup>21</sup> | 8,020,000 variants from 802 samples (10,000/sample) |
| COSMIC v103 (GenomeScreensMutant) | Aggregate somatic mutations from COSMIC genome-wide cancer screens<sup>19</sup> (no sample columns) | 3,000 (sampled from full database) |
| COSMIC v103 (NonCodingVariants) | Aggregate non-coding somatic variants from COSMIC<sup>19</sup> (no sample columns) | 3,000 (sampled from full database) |

### Datasets

Validation used a panel of real-world VCF files from 15 caller types and VCF formats, including single-sample germline (DeepVariant^9^, FreeBayes^10^, Strelka2 germline^11^, GIAB benchmark consensus^12^), paired tumor/normal somatic (GATK MuTect2^13^, Strelka2 somatic^11^, VarScan2^14^, VarDict^15^, SomaticSniper^16^), structural variant (Manta^17^ /DELLY^18^), and annotation-database VCFs (COSMIC v103^19^). Germline datasets were included alongside somatic ones because the field-mapping and allele-normalisation logic that mafsmith reimplements is exercised by the same VCF-level constructs (multi-allelic sites, FORMAT-field conventions, no-call and homozygous-reference genotypes) regardless of whether a variant is germline or somatic; germline benchmarks (GIAB) additionally provide multi-million-variant truth callsets with which we could stress-test conversion at full scale. Datasets spanned both GRCh38 and GRCh37 reference builds; full source information is provided in the Data Availability section and Table 1. For each dataset, a random subset of variants was compared field-by-field between mafsmith and vcf2maf; full-cohort validation was performed for the GIAB germline benchmarks^12^ (27.5 million variants from 7 samples), ICGC PCAWG consensus^20^ (21.6 million variants from 1,902 samples), and DepMap CCLE WGS^21^ (8 million variants from 802 samples). Beyond the datasets enumerated in Table 1, additional VCFs from the Johns Hopkins NF1 nerve sheath tumor biospecimen repository^22,23^ and a small number of caller-specific files from other NF Data Portal studies (see Data and code availability) were used ad-hoc during development to surface edge cases (for example, an Ion Torrent multi-allelic VCF that exposed an AO indexing bug, and a paired-tumor/normal VCF that exposed a `Match_Norm_Seq_Allele2` selection bug) and to confirm consistency between vcf2maf and mafsmith on caller-format permutations not represented in the primary cohorts.

### Compute cost and carbon estimation

Compute cost and carbon emissions were estimated from the wall-clock timings reported in the Performance results using AWS on-demand pricing for the c6a.4xlarge instance type ($0.612/hr, us-east-1). Power was modelled using the Cloud Carbon Footprint (CCF) methodology with AMD EPYC Milan CPU coefficients (idle: 1.04 W/vCPU; peak: 5.12 W/vCPU; DRAM: 0.392 W/GB; PUE: 1.2), giving 113.4 W at full 16-core utilisation for the mafsmith and mafsmith + fastVEP pipelines and 39.9 W for single-threaded vcf2maf. Carbon intensity was taken as 0.386 kg CO_2_e/kWh (EPA eGRID 2022^24^, SRVC subregion, Virginia, location-based). For the full annotated pipeline the same power model was applied with both mafsmith + fastVEP and vcf2maf + VEP `--vep-forks 16` assumed at 100% CPU utilisation.

## Implementation

### Overview

mafsmith is implemented in Rust and structured as a command-line tool with five subcommands: `vcf2maf` (VCF to MAF), `maf2vcf` (MAF to VCF), maf2maf (MAF re-annotation), `vcf2vcf` (VCF normalisation), and `fetch` (reference data download). The core conversion pipeline consists of: (1) optional annotation through fastVEP to produce a VCF with embedded CSQ INFO fields (CSQ is the Ensembl VEP convention for encoding per-transcript consequence data within the VCF INFO column); (2) parsing of annotated VCF records; (3) transcript selection; (4) allele normalisation; (5) genotype and depth field extraction; and (6) MAF record serialisation.

The `maf2vcf` subcommand reconstructs a standard VCF from a MAF file, recovering allele representations, multiallelic sites, genotype strings, and allele depth fields from the MAF columns. Insertion positions follow the MAF convention, with `Start_Position` equal to the VCF anchor base position. Deletions use `Start_Position` − 1 as the VCF anchor. Multi-allelic sites are reconstructed with FASTA-backed anchor bases, covering three cases: compound heterozygous tumor calls in which both tumor alleles differ from the reference; paired tumor/normal sites where the matched normal sample carries a different alt allele than the tumor; and multi-allelic deletions in which one allele is a full deletion of the reference span and another is a partial deletion sharing the same anchor base.

The `vcf2vcf` subcommand normalises a VCF: it passes through all FORMAT fields including non-PASS variants and multi-allelic ALTs, selecting the specified tumour and normal sample columns. The `maf2maf` subcommand re-annotates an existing MAF by internally converting it to VCF (`maf2vcf`), running fastVEP annotation, and converting back to MAF (`vcf2maf`).

### Annotation

When annotation is required, mafsmith invokes fastVEP^8^ with HGVS notation enabled. fastVEP produces a standard VCF with CSQ INFO fields using the same format as Ensembl VEP, allowing mafsmith to reuse the same downstream parsing logic regardless of whether annotation was performed upstream. For pre-annotated VCFs, annotation can be skipped with `--skip-annotation`. mafsmith is also compatible with the standard Ensembl VEP Perl implementation: users who prefer or require VEP can run it independently and pass the resulting annotated VCF to mafsmith with `--skip-annotation`.

### Transcript selection

mafsmith selects the canonical transcript for each variant using the same priority order as vcf2maf: (1) user-supplied custom ENST list; (2) MANE Select transcript; (3) Ensembl canonical transcript; (4) longest transcript. When multiple transcripts tie on all criteria, the first in VEP output order is used.

### Allele normalisation

VCF alleles are left-aligned and trimmed using a prefix/suffix-stripping approach. Structural variant ALT alleles (`<DEL>`, `<DUP>`, `<INV>`, BND notation) receive special handling: symbolic ALTs are mapped to their MAF representations, BND ALT strings are parsed to extract the partner chromosome and position for secondary breakpoint rows, and unrecognised symbolic ALTs (e.g. `<INS>, <CNV>`) are dropped, matching vcf2maf behaviour.

### Genotype and depth extraction

mafsmith implements vcf2maf logic for determining `Tumor_Seq_Allele1` and `Tumor_Seq_Allele2` from VCF FORMAT fields. This includes:

- **GT-based allele assignment**: GT allele indices are sorted; the minimum index determines Allele1 (REF for heterozygous, ALT for homozygous-alt).
- **Depth-based hom-alt inference**: when GT is a no-call (./.) or homozygous-reference (0/0), allele assignment falls back to AD-based VAF; VAF ≥ 0.7 is treated as homozygous-alt. This matches vcf2maf behaviour across DRAGEN^25^, MuTect2, and GVCF-style callers.
- **VAF override**: for paired tumour/normal VCFs, when GT indicates heterozygous but VAF ≥ 0.7 (suggesting caller under-calling), Allele1 is overridden to ALT, matching vcf2maf. This override is suppressed for single-sample VCFs (absent normal column).
- `--strict` **mode**: when AD arrays are shorter than the expected `1 + n_alts` length (a GATK behaviour for trimmed multi-allelic sites), `--strict` outputs. for all depth fields and suppresses depth-based allele calling, exactly matching vcf2maf. In default mode, mafsmith extracts whatever depth information is available.
- **Strelka2 somatic FORMAT fields**: Strelka2 somatic VCFs lack a GT field and use caller-specific depth fields (AU/CU/GU/TU for SNVs, TAR/TIR for indels). mafsmith extracts depth counts from these fields and infers het/hom-alt from VAF.

### Parallelism and performance

The conversion step (post-annotation) uses Rayon^26^ for parallel processing and jemalloc^27^ for improved throughput on multi-threaded runs. CSQ field parsing is deferred until a record is selected for output, reducing unnecessary work for filtered or dropped records.

## Results

### Validation

We validated mafsmith against vcf2maf across fifteen caller types and VCF formats, comparing conversion fields (`Variant_Type`, `Reference_Allele`, `Tumor_Seq_Allele1`,`Tumor_Seq_Allele2`, `Start_Position`, `End_Position`, `t_depth`, `t_ref_count`, `t_alt_count`, `n_depth, n_ref_count`, `n_alt_count`) on the same input (Table 1). For all caller types, mafsmith produced 0 conversion-field mismatches in `--strict` mode across a total of 87.2 million variants. Remaining differences between tools are restricted to `Variant_Classification` for 2–5 variants per dataset at gene-boundary regions where tools select different canonical transcripts.

To confirm that mafsmith produces equivalent output to vcf2maf when both tools use the same annotation engine, we ran end-to-end pipeline comparisons against two different Ensembl VEP releases (VEP 112 and VEP 115), each used as the shared annotation source on both sides of the comparison; the VEP 115 cache was the same one used for the timing benchmarks (Tables 5 and 8), so the equivalence and performance experiments were performed alongside one another and use a common VEP installation for the timed runs. We selected 23 representative datasets spanning all major caller types in Table 1 (20 GRCh38 datasets; 3 representative samples from the GRCh37 ICGC PCAWG consensus callset), sampling 2,000 variants per dataset, and compared MAF output from mafsmith and vcf2maf in strict comparison mode at each VEP version. `Variant_Classification` was excluded from this comparison because even with identical VEP output, mafsmith and vcf2maf.pl differ in how they handle regulatory-feature CSQ entries (ENSR-prefixed IDs): mafsmith excludes these from transcript ranking while vcf2maf.pl includes them, producing approximately 1 classification difference per 3,000 variants at gene-boundary regions; this does not affect allele-level conversion fields. Across all 23 datasets, both genome builds, and both VEP releases, mafsmith and vcf2maf produced 0 conversion-field differences, confirming that the two tools are interchangeable as drop-in replacements when using the same VEP annotation cache.

### Validation of maf2vcf, vcf2vcf, and maf2maf subcommands

We validated mafsmith’s `maf2vcf`, `vcf2vcf`, and `maf2maf` subcommands against the corresponding reference implementations from the vcf2maf Perl package (maf2vcf.pl, vcf2vcf.pl, and maf2maf.pl) across six representative datasets including GRCh38 and GRCh37 (Table 2).

**Table 2.** Validation of maf2vcf, vcf2vcf, and maf2maf subcommands against reference Perl implementations (2,000 variants per dataset sampled from datasets whose full-dataset conversion was validated in Table 1; 0 conversion differences in all cases).

| Dataset | Genome | Subcommands validated |
| --- | --- | --- |
| SEQC2 HCC1395, GATK MuTect2 | GRCh38 | `maf2vcf`, `vcf2vcf`, `maf2maf` |
| SEQC2 HCC1395, Strelka2 somatic | GRCh38 | `maf2vcf`, `vcf2vcf` |
| SEQC2 HCC1395, SomaticSniper | GRCh38 | `maf2vcf`, `vcf2vcf` |
| GIAB HG008, GATK MuTect2 | GRCh38 | `maf2vcf`, `vcf2vcf`, `maf2maf` |
| GIAB HG001 germline benchmark | GRCh38 | `maf2vcf`, `vcf2vcf` |
| PCAWG consensus (0009b464) | GRCh37 | `maf2vcf`, `vcf2vcf`, `maf2maf` |

For `maf2vcf` and `vcf2vcf`, comparisons used 2,000 variants per dataset. Input MAFs for the `maf2vcf` comparison were generated by running `mafsmith` `vcf2maf` `--skip-annotation` on the same VCFs used in Table 1, ensuring that both `maf2vcf` tools (mafsmith and maf2vcf.pl) received byte-identical inputs; this isolates any output differences to `maf2vcf` logic rather than compounding upstream vcf2maf differences. For `vcf2vcf`, both tools processed the same input VCF directly. For maf2maf, comparison used 2,000 variants per dataset with VEP 112 annotation for the maf2maf.pl reference and fastVEP for mafsmith; `Variant_Classification` was excluded from comparison because the two annotation engines may use different GFF3 gene model releases, producing transcript-selection differences independent of conversion logic.

Achieving concordance required resolving several non-obvious behaviours in the reference implementations. For `maf2vcf`, key fixes included: correctly applying the MAF insertion position convention (where `Start_Position` is the anchor base, matching the VCF POS, not one position upstream); reconstructing multi-allelic deletions where the effective alt is a full deletion and TSA1 encodes a partial deletion (both anchored to the same VCF POS − 1); and correctly assigning GT=1/1 (homozygous alt) for deletions where TSA1 and TSA2 are both “-”. For `vcf2vcf`, the reference implementation passes non-PASS variants and preserves all multi-allelic ALT alleles; mafsmith was updated to match this behaviour. For `maf2maf`, mafsmith was updated to output vcf2maf-compatible depth field defaults (total depth → “0”, allele counts → “.”) when a sample column is present but no depth FORMAT fields exist.

### Performance

We benchmarked mafsmith against vcf2maf on the conversion step in isolation, passing the same raw VCF to each tool (`mafsmith` `--skip-annotation` and vcf2maf --inhibit-vep) without any prior annotation. Benchmarks were run on an AWS c6a.4xlarge instance (AMD EPYC 7R13, 16 vCPU, 30 GiB RAM) using seven GIAB NIST v4.2.1 GRCh38 benchmark VCFs (HG001–HG007; 3.84–4.05 million variants per file; 27.5 million variants total). To distinguish algorithmic speedup from parallelism, mafsmith was timed at both 1 core (`RAYON_NUM_THREADS=1`, 3 runs per sample) and 16 cores (default Rayon, 1 run per sample); vcf2maf is single-threaded. Per-sample timings are reported in Table 3.

**Table 3.** Conversion-only benchmark: mafsmith vs vcf2maf on GIAB NIST v4.2.1 GRCh38.

| Sample | Variants | mafsmith 1-core (s) | mafsmith 16-core (s) | vcf2maf (s) | Speedup (1-core) | Speedup (16-core) |
| --- | --- | --- | --- | --- | --- | --- |
| HG001 (NA12878) | 3,893,341 | 11.6 | 7.40 | 550 | 47.5× | 74.3× |
| HG002 (NA24385) | 4,048,342 | 12.3 | 7.40 | 580 | 47.1× | 78.3× |
| HG003 (NA24149) | 4,000,097 | 12.1 | 7.19 | 574 | 47.5× | 79.7× |
| HG004 (NA24143) | 4,031,346 | 12.0 | 7.05 | 574 | 47.8× | 81.3× |
| HG005 (NA24631) | 3,856,856 | 12.0 | 7.21 | 558 | 46.6× | 77.4× |
| HG006<br>(NA24694) | 3,839,315 | 11.1 | 6.67 | 538 | 48.6× | 80.7× |
| HG007<br>(NA24695) | 3,859,704 | 11.3 | 6.42 | 540 | 48.0× | 84.1× |
| **Mean**<br>**± SD** |  | **11.8 ± 0.5 s** | **7.05 ± 0.37 s** | **559 ± 17 s** | **47.6× ± 0.6×** | **79.4× ± 3.1×** |

On a single core, mafsmith achieved a mean throughput of 334,802 variants/s (a 47.6-fold speedup over vcf2maf; range 46.6–48.6×). The performance advantage is therefore primarily algorithmic rather than a product of parallelism. With all 16 cores, throughput increased to 558,914 variants/s (79.4-fold speedup, range 74.3–84.1×), a 1.67× parallel scaling factor. Both speedups were consistent across samples (coefficient of variation: 1.3% and 3.9%, respectively).

To confirm that speedups generalise to paired tumor/normal somatic VCFs, we benchmarked mafsmith on five additional datasets: MuTect2 and Strelka2 VCFs from the GIAB HG008 somatic benchmark (NYGC pipeline, GRCh38; HG008-T / HG008-N), and MuTect2 and Strelka VCFs from the SEQC2 WGS somatic dataset (HCC1395 / HCC1395BL). All five VCFs carried paired tumor and normal sample columns. Per-sample timings are reported in Table 4.

**Table 4.** Somatic tumor/normal benchmark: mafsmith vs vcf2maf.

| Dataset / Caller | Variants | mafsmith 1-core (s) | mafsmith 16-core (s) | vcf2maf (s) | Speedup (1-core) | Speedup (16-core) |
| --- | --- | --- | --- | --- | --- | --- |
| GIAB HG008 / MuTect2 | 277,645 | 1.29 | 0.779 | 42.4 | 32.8× | 54.4× |
| GIAB HG008 / Strelka2 SNV | 1,562,847 | 5.27 | 2.96 | 246 | 46.6× | 83.1× |
| GIAB HG008 / Strelka2 INDEL | 293,719 | 1.24 | 0.684 | 44.7 | 36.1× | 65.3× |
| SEQC2 HCC1395 / MuTect2 | 271,945 | 1.18 | 0.631 | 38.8 | 32.9× | 61.4× |
| SEQC2 HCC1395 / Strelka | 2,191,720 | 7.30 | 4.15 | 346 | 47.3× | 83.3× |
| **Mean** |  |  |  |  | **39.1×** | **69.5×** |

mafsmith achieved a mean single-core speedup of **39.1×** (range 32.8–47.3×) and a 16-core speedup of **69.5×** (range 54.4–83.3×) across paired tumor/normal VCFs. The lower bound of the range reflects MuTect2 VCFs, which carry larger per-variant INFO fields (TLOD, NLOD, per-allele annotations) that increase per-line parsing cost for both tools. Note also that vcf2maf does not accept gzip-compressed input and requires decompression before processing; mafsmith reads gzip natively, a further practical advantage not captured in these timings.

To quantify the full annotation pipeline speedup, we re-ran all five datasets with annotation enabled. We used VEP with `--vep-forks 16` to match fastVEP’s 16-core Rayon parallelism. Table 5 reports the symmetric comparison.

**Table 5.** Full annotated pipeline benchmark: mafsmith + fastVEP vs. vcf2maf + VEP 115 (symmetric 16-core comparison).

| Dataset / Caller | Variants | mafsmith 1-core (s) | mafsmith 16-core (s) | vcf2maf + VEP (s) | Speedup (1-core) | Speedup (16-core) |
| --- | --- | --- | --- | --- | --- | --- |
| GIAB HG008 / MuTect2 | 277,645 | 25.8 | 11.6 | 460 | 17.8× | 39.6× |
| GIAB HG008 / Strelka2 SNV | 1,562,847 | 94.0 | 30.9 | 2,850 | 30.3× | 92.4× |
| GIAB HG008 / Strelka2 INDEL | 293,719 | 25.7 | 11.6 | 613 | 23.8× | 53.0× |
| SEQC2 HCC1395 / MuTect2 | 271,945 | 20.0 | 10.9 | 446 | 22.2× | 40.9× |
| SEQC2 HCC1395 / Strelka | 2,191,720 | 129 | 41.0 | 4,170 | 32.4× | 101.7× |
| **Mean** |  |  |  |  | **25.3×** | **65.5×** |

Even in the symmetric 16-core comparison, the full mafsmith + fastVEP pipeline achieved a mean single-core speedup of **25.3×** (range 17.8–32.4×) and a 16-core speedup of **65.5×** (range 39.6–101.7×).

Annotation engine choice is orthogonal to the conversion logic that is the subject of this work, and mafsmith accepts pre-annotated input from either fastVEP or the Ensembl VEP Perl implementation. We note in passing that fastVEP and VEP 115 produced consequence-level agreement for 87.6% of variants in a 1,000-variant comparison on the HG008 MuTect2 VCF; the remainder was driven by gene-model version drift between the two installations rather than by differences in consequence-classification logic.

### Compute cost and carbon savings

Despite drawing more instantaneous power at full 16-core utilisation (113.4 W vs. 39.9 W for single-threaded vcf2maf), the far shorter wall-clock time gave mafsmith substantially lower total energy and carbon per sample on the conversion-only benchmark (GIAB NIST v4.2.1 GRCh38, mean across HG001–HG007): 0.22 Wh vs. 6.20 Wh, and 0.086 g vs. 2.393 g CO_2_e, for a per-sample saving of 5.97 Wh and 2.31 g CO_2_e (Table 6). At cohort scale these per-sample savings compound to 23.1 kg CO_2_e and $938 in compute for 10,000 samples (Table 7).

**Table 6.** Estimated compute cost and carbon emissions per sample (GIAB NIST v4.2.1 GRCh38, conversion step only; mean across HG001–HG007; c6a.4xlarge, us-east-1).

| Metric | mafsmith (16-core) | vcf2maf | Saved per sample |
| --- | --- | --- | --- |
| Wall-clock time | 7.1 s | 559.1 s | 552.0 s |
| Instance cost | \$0.00120 | \$0.09504 | **\$0.094** |
| Estimated power draw | 113.4 W | 39.9 W | - |
| Energy consumed | 0.22 Wh | 6.20 Wh | **5.97 Wh** |
| CO<sub>2</sub>e (location-based) | 0.086 g | 2.393 g | **2.31 g** |

**Table 7.** Estimated compute cost and carbon savings at scale (conversion step only; c6a.4xlarge on-demand, us-east-1; estimated per sample assuming approximately 3.93 million variants per sample, the mean across GIAB HG001–HG007).

| Cohort size | Compute cost saved | CO<sub>2</sub>e avoided | Wall-clock time saved |
| --- | --- | --- | --- |
| 100 samples | \$9.38 | 0.23 kg | 15.3 hr |
| 1,000 samples | \$93.84 | 2.31 kg | 153 hr |
| 10,000 samples | \$938 | 23.1 kg | 1,533 hr |
| 100,000 samples | \$9,384 | 231 kg | 15,334 hr |
| 1,000,000 samples | \$93,844 | 2,307 kg | 153,340 hr |

For the full annotated pipeline, mean time savings of 1,686 s per run corresponded to $0.287 in compute cost and 20.5 g CO_2_e (Table 8). Per million variants, the annotated pipeline saves approximately 21.8 g CO_2_e, compared with 2.3 g CO_2_e for the conversion step alone (Table 6), reflecting the dominance of the annotation bottleneck. Projected savings at cohort scale are shown in Table 9.

**Table 8.** Full annotated pipeline compute cost and carbon savings per run (mafsmith + fastVEP 16-core vs. vcf2maf + VEP 115 --vep-forks 16; c6a.4xlarge, us-east-1).

| Dataset | Variants | Time saved (s) | Cost saved | Energy saved (Wh) | CO<sub>2</sub>e saved (g) |
| --- | --- | --- | --- | --- | --- |
| GIAB HG008, MuTect2 | 277,645 | 448 | \$0.076 | 14.1 | 5.45 |
| GIAB HG008, Strelka2 SNV | 1,562,847 | 2,821 | \$0.480 | 88.8 | 34.3 |
| GIAB HG008, Strelka2 INDEL | 293,719 | 602 | \$0.102 | 18.9 | 7.31 |
| SEQC2 HCC1395, MuTect2 | 271,945 | 435 | \$0.074 | 13.7 | 5.28 |
| SEQC2 HCC1395, Strelka | 2,191,720 | 4,126 | \$0.701 | 129.9 | 50.2 |

**Table 9.** Projected full-pipeline compute cost and carbon savings at scale (mean per run across five WGS somatic datasets, ∼920K variants; c6a.4xlarge on-demand, us-east-1).

| Cohort size | Compute cost saved | CO<sub>2</sub>e avoided | Wall-clock time saved |
| --- | --- | --- | --- |
| 100 runs | \$28.70 | 2.05 kg | 46.8 hr |
| 1,000 runs | \$287 | 20.5 kg | 468 hr |
| 10,000 runs | \$2,870 | 205 kg | 4,683 hr |
| 100,000 runs | \$28,700 | 2,050 kg | 46,833 hr |
| 1,000,000 runs | \$287,000 | 20,490 kg | 468,333 hr |

## Discussion

The primary motivation for mafsmith is throughput: converting thousands of VCFs in large cancer genomics cohorts places substantial demands on compute infrastructure when using vcf2maf, and the per-sample annotation step is a major bottleneck. Combining fastVEP and mafsmith achieves substantially faster conversion. The MAF conversion logic (field population, allele assignment, consequence mapping) maintains field-for-field agreement with vcf2maf on all evaluated datasets, while annotation-level concordance with VEP 115 depends on the Ensembl gene model version used by fastVEP. The single-core benchmark (47.6× speedup) confirms that the performance advantage is primarily algorithmic, with parallelism providing an additional 1.67× with 16 virtual cores.

Achieving full concordance with vcf2maf required identification and implementation of a number of non-obvious behaviours accumulated over years of production use. Examples include: the treatment of absent normal samples vs. no-call normal GTs in homozygous-alt inference; the VAF-based Allele1 override and its interaction with single-sample vs. paired VCF configurations; the handling of truncated AD arrays from GATK multi-allelic calling; consequence severity ranking for multi-consequence VEP annotations; and the representation of structural variant secondary breakpoint rows.

Several known differences remain. In default mode (without `--strict`), mafsmith extracts partial depth counts when AD arrays are truncated, which may be more informative than outputting missing values. mafsmith populates SV secondary breakpoint rows with the partner chromosome and position, whereas vcf2maf leaves these fields blank. These intentional differences are documented and `--strict` mode is provided for workflows requiring exact vcf2maf compatibility. mafsmith currently uses a prefix/suffix-stripping approach for allele normalisation rather than FASTA-guided left-alignment. For most variant types this produces identical results to vcf2maf, but complex indels at homopolymer runs or short tandem repeats may differ. Full FASTA-based normalisation is planned for a future release.

Faster conversion also translates into measurable reductions in compute energy and carbon emissions: at the conversion step alone mafsmith saved approximately 2.3 g CO_2_e per sample, which compounds to ∼23 kg CO_2_e for a 10,000-sample cohort, and the full annotated pipeline (mafsmith + fastVEP vs. vcf2maf + VEP 115) saves a further ∼20.5 g CO_2_e per run on average.

mafsmith is a complete, self-contained rewrite of vcf2maf in Rust, designed as a drop-in replacement. It produces field-for-field identical MAF output across fifteen distinct caller types and VCF formats, spanning germline, somatic, structural variant, and annotation-database VCFs, while achieving 47.6-fold faster conversion on a single core (range 46.6–48.6×) and 79.4-fold faster on 16 cores (range 74.3–84.1×) across seven GIAB reference samples totalling 27.5 million variants, making VCF-to-MAF conversion faster and more cost-efficient.

## Data and code availability

mafsmith source code and release binaries are available at github.com/nf-osi/mafsmith, together with the validation, benchmarking, and compute-cost scripts used to produce every table and figure in this paper; the `results/README`.md file in the repository gives a per-table mapping from each manuscript table to its source data and generating script.

Validation VCF datasets are available from the sources listed below. Validation and benchmark VCF datasets used in this work:

- **NF Data Portal**^33^: Validation and refinement VCFs contributed by the following NF Hub studies:
  – Preclinical NF1-MPNST platform^32^ (project syn11638893): syn21296193, syn21296175, syn21296184 (Manta/DELLY somatic SV).
  – Cutaneous neurofibroma mechanism study^28^ (project syn4939910): syn31624525 (MuTect2), syn31624535 (FreeBayes), syn31624545 (DeepVariant), syn31624637 / syn31624939 (Strelka2 germline), syn31625234 (additional variant call used during refinement).
  – Cutaneous Neurofibroma Data Resource^31^ (project syn4984604): syn6039268 (VarDict), syn68172710 (Strelka2 somatic indels), syn6840402 (VarScan2). Additional paired-T/N and tumor-only VCFs from this resource were used during development to surface and fix bugs in multi-allelic depth handling: syn5553155, syn5553158, syn5555584, syn5614682.
  – NF Cell Line Compound Screens^34^ (project syn11817821): syn20443868 (Ion Torrent multi-allelic VCF used to surface and fix an AO indexing bug).
  – Genetic Studies of Neurofibromatosis^30^ (project syn11374339): syn29349669 (additional Strelka2 germline variants file).
  – Johns Hopkins NF1 nerve sheath tumor biospecimen repository^22,23^ (project syn4939902): additional VCFs used ad-hoc during development to identify edge cases and validate consistency between vcf2maf and mafsmith. All files above are accessible through the NF Data Portal at nf.synapse.org.
- **GIAB HG008 somatic (NYGC pipeline, GRCh38)**^12^: Paired tumor/normal (HG008-T / HG008-N) MuTect2 and Strelka2 VCFs. Available from the GIAB HG008 NYGC-somatic-pipeline FTP directory.
- **SEQC2 WGS somatic (HCC1395 / HCC1395BL, GRCh38)**^29^: Paired tumor/normal MuTect2, Strelka2, and SomaticSniper VCFs from the FDA Sequencing Quality Control Phase II (SEQC2) Somatic Mutation Working Group. Available from the SEQC2 Somatic Mutation WG FTP directory.
- **COLO829 somatic SV truth set (hg38)**^35^: Somatic structural variant truth set for the COLO829 melanoma cell line. Lift-over to GRCh38: `truthset_somaticSVs_COLO829_hg38lifted.vcf`. Available on Zenodo record 7515830.
- **GIAB germline benchmarks (HG001–HG007, GRCh38)**^12^: NIST GIAB v4.2.1 benchmark VCFs used for conversion-speed benchmarking and validation. Available from the GIAB release FTP directory.
- **COSMIC v103 (GRCh38)**^19^: Genome Screens Mutant (Normal) and Non-Coding Variants VCFs used for validation of annotation-database VCF format handling. Available under the COSMIC licence from the COSMIC download portal.
- **ICGC PCAWG cell-line VCFs (GRCh37)**^20^: DKFZ SNV/MNV somatic VCFs for HCC1143 and HCC1954 cell lines from the ICGC Pan-Cancer Analysis of Whole Genomes (PCAWG) open data release. Available through the ICGC-ARGO open-access S3 endpoint (bucket: `icgc25k-open;` no sign request required for open-tier data).

## Acknowledgements

mafsmith builds on the design, field conventions, and years of accumulated edge-case handling embodied in vcf2maf. We are grateful to Cyriac Kandoth and other vcf2maf contributors for their sustained work developing and maintaining vcf2maf without which this rewrite would not have been possible. This article was drafted with the assistance of large language models (Anthropic Claude Sonnet 4.6), with the author reviewing, editing, and verifying all content.

